# Transcriptome signature of cell viability predicts drug response and drug interaction for Tuberculosis

**DOI:** 10.1101/2021.02.09.430468

**Authors:** Vivek Srinivas, Rene A. Ruiz, Min Pan, Selva Rupa Christinal Immanuel, Eliza J.R. Peterson, Nitin S. Baliga

## Abstract

The treatment of tuberculosis (TB), which kills 1.8 million each year, remains difficult, especially with the emergence of multidrug resistant strains of *Mycobacterium tuberculosis* (Mtb). While there is an urgent need for new drug regimens to treat TB, the process of drug evaluation is slow and inefficient owing to the slow growth rate of the pathogen, the complexity of performing bacteriologic assays in a high-containment facility, and the context-dependent variability in drug sensitivity of the pathogen. Here, we report the development of “DRonA” and “MLSynergy”, algorithms to perform rapid drug response assays and predict response of Mtb to novel drug combinations. Using a novel transcriptome signature for cell viability, DRonA accurately detects bacterial killing by diverse mechanisms in broth culture, macrophage infection and patient sputum, providing an efficient, and more sensitive alternative to time- and resource-intensive bacteriologic assays. Further, MLSynergy builds on DRonA to predict novel synergistic and antagonistic multi-drug combinations using transcriptomes of Mtb treated with single drugs. Together DRonA and MLSynergy represent a generalizable framework for rapid monitoring of drug effects in host-relevant contexts and accelerate the discovery of efficacious high-order drug combinations.

## Introduction

New treatment regimens containing multiple drugs are needed to achieve rapid and complete clearance of *Mycobacterium tuberculosis* (Mtb), the causative agent of tuberculosis (TB). However, the discovery of effective multidrug combinations is a challenging endeavor, burdened by the enormous number of testable combinations (e.g., a collection of 1,000 compounds yields ∼500,000 pairwise combinations, and exponentially larger numbers of higher-order combinations). Multicomponent drug discovery is particularly challenging for Mtb, a slow-growing pathogen that is capable of generating phenotypically heterogeneous subpopulations. These phenotypically diverse subpopulations allow Mtb to persist and survive the variable conditions encountered during infection as well as thwart drug treatment. Because of drug tolerant subpopulations within the host, a large proportion of drug regimens that are effective in killing Mtb *in vitro* are futile in patients^1–3^. Suffice it to say, new approaches are needed to reduce the search space and prioritize combinations for experimental testing while also taking into account the host context and different subpopulations of Mtb.

Another challenge in the development of new antitubercular drug regimens is the reliance on growth assays to monitor treatment response. Current methods to monitor treatment response include counting of colony forming units (CFUs) on solid agar plates, and measuring the time it takes for a sample in liquid culture to become culture positive for Mtb, in what is termed time to positivity (TTP) assay. Both CFU counting and TTP have their drawbacks including loss of sensitivity, vulnerability to contamination and lengthy time to measure results. Furthermore, culture on solid media or in liquid media requires actual growth, which limits the detection of mycobacterial subpopulations that are viable but not actively growing^4^. Instead, profiling 16S ribosomal RNA as a proxy for Mtb load in sputum is emerging as a more sensitive technique that addresses the shortcomings of growth-based assays^5,6^. This is a promising development because information in RNA can be amplified using technologies like probe-capture and PCR to develop highly-sensitive methods for investigating the drug response of Mtb, especially from patient samples. Furthermore, genome-wide expression studies of Mtb from broth, sputum and *in vivo* infections have been used to uncover physiologic states and transcriptional mechanisms of drug tolerance^7–9^. Here, we have investigated if transcriptome profiling of Mtb can report on the effect of drug treatment, and whether this information can also enable *in silico* identification of drug combinations that are likely to have synergistic or antagonistic effects on the pathogen.

We report the development of a framework of two algorithms “DRonA’’ and “MLSynergy” that can use transcriptomes to predict Mtb’s response to drug treatment and classify 2- and 3-drug combinations based on likelihood of synergistic or antagonistic action on Mtb. DRonA is a machine learning algorithm that was trained on publicly available transcriptomes of Mtb cultured in diverse conditions (with and without perturbation) to detect a gene signature for loss of Mtb viability. Using drug-induced transcriptional changes, DRonA calculates a cell viability score (CVS), which distinguishes the extent of a drug’s bacteriostatic or bactericidal activity on Mtb. We demonstrate that DRonA accurately detects within the transcriptome profile of drug-treated Mtb, evidence for loss of bacterial viability, regardless of the drug’s mechanism of killing. Furthermore, DRonA was equally accurate in determining drug-induced viability reduction in Mtb from broth culture, macrophage infection, and patient sputum. Finally, MLSynergy uses transcriptomes from single-drug treatment to prediction the interaction of drugs in combination. Using the ratio of the expected CVS (based on CVS of individual drugs) and the predicted CVS (based on the inferred multi-drug transcriptome) calculated by DRonA, MLSynergy can distinguish between synergistic and antagonistic combinations. We demonstrate that MLSynergy accurately classified experimentally-determined synergistic and antagonistic combinations. Thus, DRonA/MLSynergy framework can accelerate antitubercular drug discovery by reducing the reliance on growth-based treatment response assays and guiding the experimental assessment of novel drug combinations.

## Results

### <u>D</u>rug <u>R</u>esp<u>on</u>se <u>A</u>ssayer (DRonA) detects signatures for loss of viability within transcriptomes of Mtb irrespective of mechanism of killing

To investigate whether Mtb viability can be deciphered from its transcriptome state, we sought to define a classifier that could accurately identify transcriptomes of viable Mtb. We hypothesized that the degree of deviation of a transcriptome profile from classifier-defined viable transcriptomes could accurately predict the loss of viability of Mtb cells. Further, we hypothesized that the loss of viability would be agnostic of the inhibitory effect, making it possible to predict drug-mediated killing regardless of the drug’s mechanism of action. We trained a single class support vector machine (SC-SVM) model on a compendium of transcriptomes of Mtb from diverse growth conditions. In order to train the SC-SVM classifier, we compiled a compendium of 3,151 transcriptomes available in the Gene Expression Omnibus (GEO) from 173 studies. These studies used microarray and RNA-sequencing (RNA-seq) to assess gene expression changes in Mtb from various growth medium compositions, culture conditions, and drug treatment (**File S1**). Batch-effects and platform-specific bias across the transcriptome profiles were corrected with rank normalization and then used as input to train the SC-SVM classifier (**Figure 1**). Specifically, training of the SC-SVM was seeded with 24 transcriptomes of Mtb cultures from optimal growth conditions (mid-log phase of growth in 7H9 medium, 37° C, shaking,) as “viable” transcriptomes. New datasets were iteratively added to the set of “viable” transcriptomes, with simultaneous evaluation of the SC-SVM classifier performance in differentiating the viable transcriptomes from transcriptomes of Mtb treated with drugs (193 transcriptomes, 17 drugs). The training process was iterated until addition of new samples caused a drop in the performance of the classifier (**Figure S1**). The final classifier was trained on 994 transcriptomes (**File S1**) of Mtb from diverse growth conditions, including log phase, vehicular control samples, nutrient starvation, low pH, hypoxia, and intracellular growth. As such, the SC-SVM model, identified Mtb transcriptomes from slow growing (i.e., dormancy-inducing), but viable conditions. In contrast, the excluded transcriptomes (total 1,940) (**File S1**) were from conditions lethal to Mtb (e.g., drug-treated, detergent-treated, heat-treated etc.) and from genetically perturbed strains of Mtb (e.g. *phoP, espR, mihF* etc. mutants).

**Figure 1.**
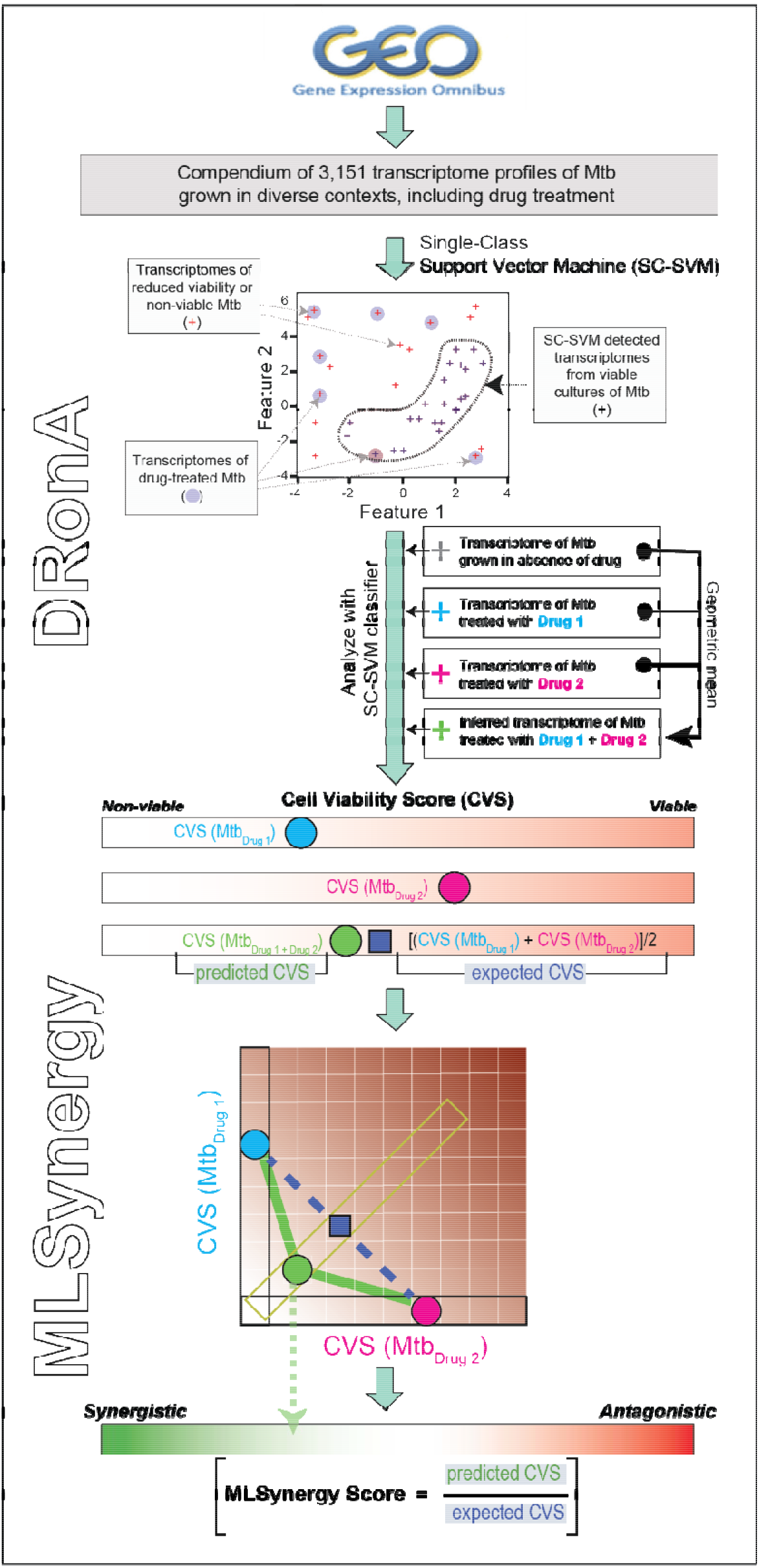
Overview schematic of DRonA/MLSynergy framework. DRonA is a single-class support vector machine classifier (SC-SVM) that was trained on transcriptomes from viable Mtb cultures. DRonA was trained through an iterative process to define a region in the hyperplane which classifies transcriptomes from Mtb grown in varying growth conditions as viable and distinguishes them from non-viable transcriptomes (i.e., drug treated at >MIC_50_ concentration). DRonA takes transcriptomes as input and outputs a cell viability score (CVS), which is the empirical distance from the viable class and indicative of efficacious drug treatment. Using an inferred transcriptome of a drug combination from single-drug transcriptomes, MLSynergy predicts the outcome of the drug interaction. MLSynergy uses the Loewe additivity principle and calculates the ratio of predicted CVS to expected CVS to determine synergy or antagony for drug combinations.

The linear SC-SVM classifier, named **D**rug **R**esp**on**se **A**ssayer (DRonA), can take as input transcriptomes of Mtb to calculate a Cell Viability Score (CVS, see methods for details). The calculated CVS is proportional to the deviation of a given transcriptome from the lower limit of the classifier defined viable transcriptome space. This lower limit is set as the cell viability threshold (cell viability threshold = -3.5e^10^), below which a CVS indicates a transcriptome signature of nonviable Mtb. Using an independent compendium of 72 transcriptomes generated for this study (**File S2**), we ascertained that the CVS scoring scheme of DRonA accurately classified as “viable” (i.e., with a CVS > -3.5e^10^) all 27 transcriptomes of Mtb grown in 7H9 medium in the absence of drugs. By contrast, DRonA predicted loss of viability (CVS < -3.5e^10^) from transcriptomes of Mtb cultures treated for 72 hours in 7H9 growth medium with each of 7 frontline TB drugs at ≥MIC_50_ concentration (*p*-value < 0.001, **Figure 2A**). As expected, pyrazinamide treatment at 3.0 mg/mL was not predicted to reduce the viability of Mtb^10^. Next, we tested the performance of DRonA in predicting Mtb viability within an intracellular host context, using as input 39 transcriptomes of Mtb from naive, lipopolysaccharide (LPS)-activated and drug-treated infected macrophages of J774A.1 lineage (**File S2**)^11^. Again, DRonA correctly classified the transcriptomes from untreated Mtb as viable and the drug-treated transcriptomes as non-viable. Moreover, DRonA detected the known increase in the intracellular efficacy of pyrazinamide, isoniazid and ethambutol^12,13^ and also the decreased efficacy of rifampicin^14^ in killing Mtb within macrophages (**Figure 2B**). DRonA also detected a loss in the viability of Mtb within interferon-gamma activated macrophages upon LPS treatment^15^. Together this demonstrates that DRonA was able to identify non-viable transcriptomes, irrespective of the context and underlying mechanism of killing (i.e., whether immune- or drug-induced). Finally, we tested the performance of DRonA in predicting drug response within TB patients, using as input 16 transcript profiles of Mtb from sputum of 8 patients at the start of and after 7 or 14 days of successful TB treatment with isoniazid (H), rifampicin (R), pyrazinamide (Z), and ethambutol (E)^7^. DRonA efficiently differentiated cell viability from the Mtb transcriptomes collected from patients on day 0 from transcriptomes collected on day 7 or 14 of drug treatment (*p*-value < 0.01) (**Figure 2C**), demonstrating that DRoNA can detect drug treatment response from bacterial RNA in patient sputum.

**Figure 2.**
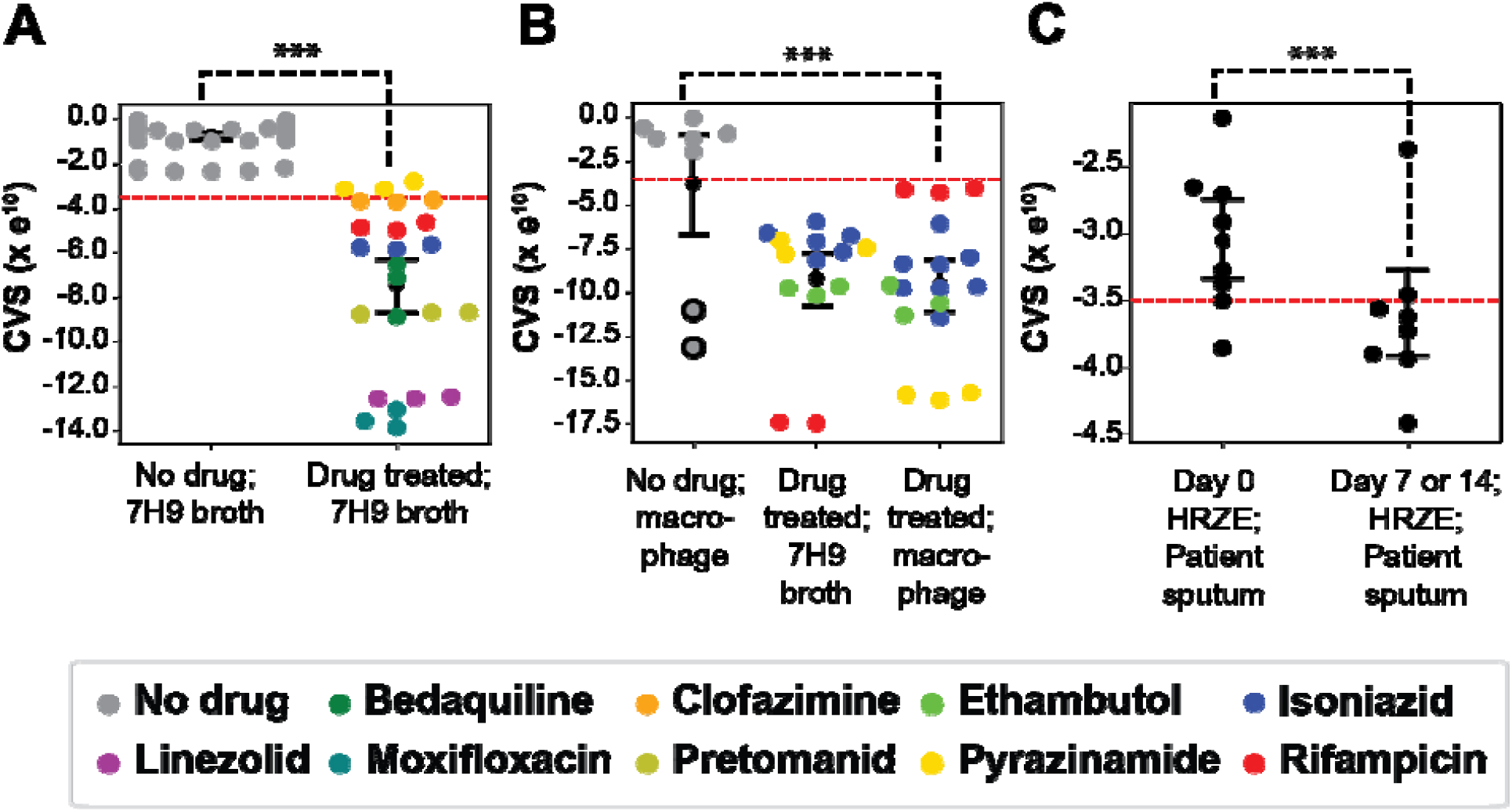
DRonA-generated CVSs for transcriptomes of Mtb sourced from broth culture, macrophage infection and patient sputum. (**A**) CVSs for transcriptomes of Mtb cultures grown in 7H9-rich media with or without drug treatment for 72 h. (**B**) CVSs for transcriptomes of Mtb cultured in 7H9 broth with drug treatment for 24 h and macrophage with or without drug treatment for 24 h. Circles with black border indicate transcriptomes from interferon gamma activated macrophages with lipopolysaccharide treatment. **C**) CVSs for transcriptomes of Mtb in patient sputum collected at the start and end of 7 or 14 day chemotherapy with HRZE; isoniazid (H), rifampicin (R), pyrazinamide (Z), and ethambutol (E). Red -dashed line is the cell viability threshold (−3.5e^10^), below which the samples are considered to be non-viable. Black dot and error bars indicate the mean and standard deviation away from the mean. Statistical significance (black dashed line) was calculated as *p*-value with Student’s T-test. ***: *p*-value < 0.001.

### DRonA can estimate the decline in CFUs upon drug treatment

We next tested whether the CVS was proportional to the magnitude of drug effects based on CFU assessment. We compared DRonA-generated CVSs with the relative CFUs observed after Mtb was treated for 24 and 72 hours with 7 frontline TB drugs at various concentrations (**Table S2)**. The CVS scores calculated from transcriptomes of both untreated Mtb cultures and those treated with drugs at <MIC_50_ concentrations were higher than the viability threshold. Although, the inferred CVS from cultures treated with <MIC_50_ drug was less than the CVS of untreated cultures (difference in average = -3.5 × e^10^, *p*-value < 0.01), indicating a moderate loss of viability. In contrast, the CVS scores calculated from transcriptomes of Mtb cultures treated with ≥MIC_50_ concentration of drug were consistently below the viability threshold. Furthermore, the reduction in CVS was proportional to decrease in CFU for most drugs, with the exception of bedaquiline (**Figure 3**). It is known that bedaquiline kills Mtb relatively slowly compared to other frontline drugs and could explain the discord between CFU and CVS within 72 h of treatment^16–20^. It is remarkable that despite the slow bactericidal activity of bedaquiline, its lethal effect was captured in transcriptome changes at a significantly earlier timepoint and we still observed a significant correlation between relative CFUs and CVS across all drug treatments (r = -0.9, *p*-value < 0.001). A notable advantage of using DRonA is that the CVS scores determined via mRNA signatures represents a comprehensive assay of drug effects on the entire Mtb population, unlike survival assessment via CFU counting, which is limited to only those subpopulations that are able to form colonies on agar plates.

**Figure 3.**
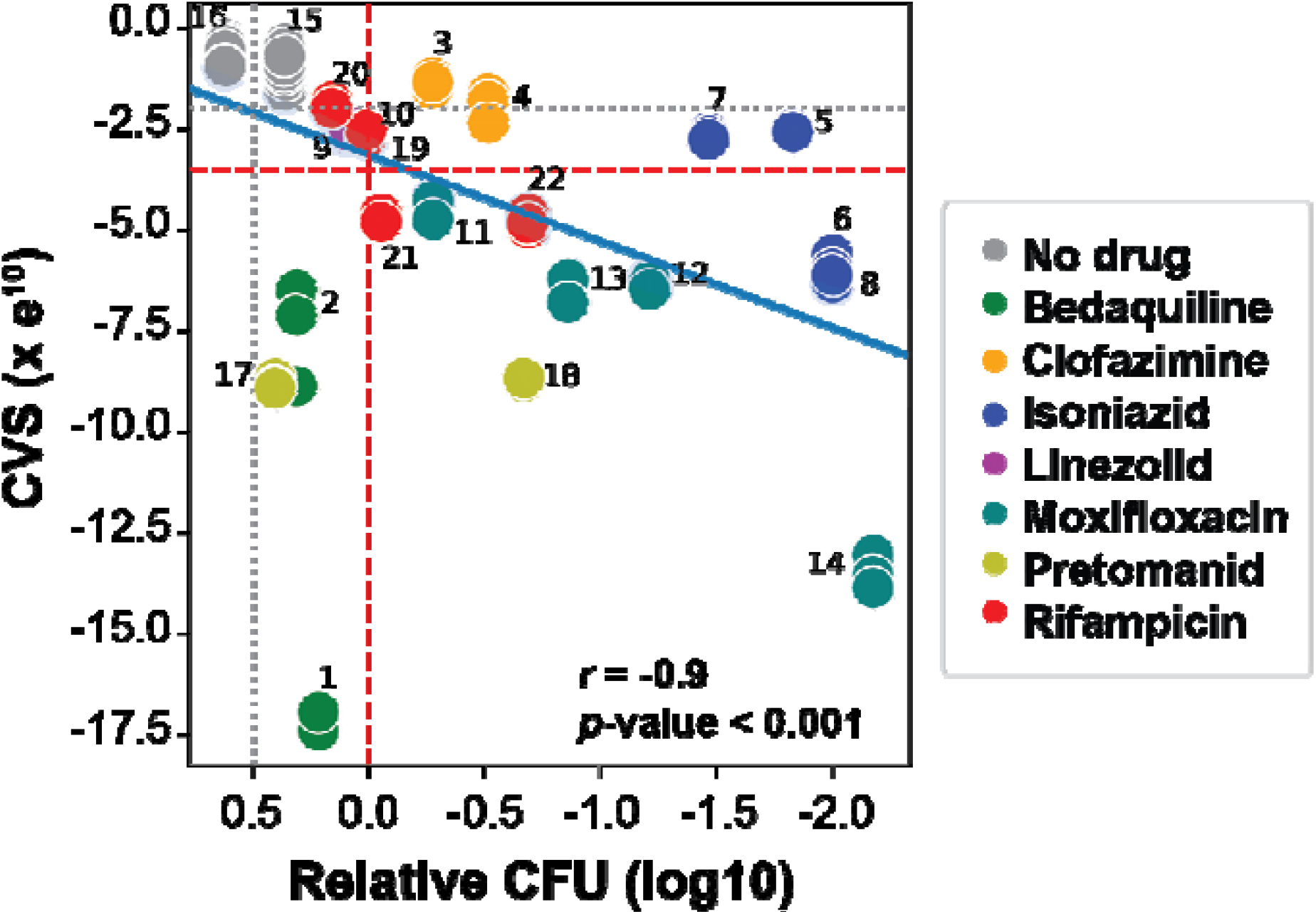
Correlation between CVS and relative CFU with and without drug treatment. Relative CFU was calculated in relation to 0 h (prior to drug or vehicle control treatment). Numbers associated with the points indicate specific drug treatment time and concentrations found in **Table S2**. Solid blue line denotes the Pearson’s correlation between CVS and relative CFU. Significance was calculated as the average correlation coefficient, *r*, from 100 iterations performed with 70% randomly selected data. Black dotted line denotes 50% growth inhibition from drug treatments and its corresponding CVS threshold (−2.25e^10^). Dashed red line indicates bactericidal activity and its corresponding CVS threshold (−3.5e^10^).

### DRonA accurately predicts synergistic and antagonistic drug combinations from transcriptomes of single drug-treated Mtb

Given that DRonA can detect Mtb’s response to drug treatment from gene expression data, we investigated if DRonA could be used to accelerate multicomponent drug discovery by predicting outcome of drug interactions from single drug-treated transcriptomes. To do this, we developed an approach to infer the transcriptomes of multi-drug treatments. Specifically, we calculated the geometric mean of two (or three) transcriptomes obtained from Mtb treated with single drugs and then predicted the CVS of the inferred drug combination transcriptome using DRonA. Using this method to infer the transcriptomes of drugs in combination and DRonA to predict their CVS, we developed a parametric method, “MLSynergy”, to predict the interaction outcome of the two- and three-drug combinations. MLSynergy predicts the synergy or antagony of multi-drug combinations based on the Loewe additivity principle^21^ by calculating the ratio of predicted CVS to expected CVS. For example, the predicted CVS of the antagonistic combination linezolid and moxifloxacin^22,23^ is greater than the expected CVS and lies above the additive plane (**Figure 4A**), whereas the predicted CVS of the synergistic combination linezolid and POA is less than the expected CVS and lies below the additive plane (**Figure 4B**). Finally, the predicted CVS of linezolid with itself is the same as the expected CVS and lies on the additive plane, consistent with the Loewe additivity principle which states that a drug in combination with itself is additive in interaction^21^ (**Figure 4C**). As such, an MLSynergy score <1 predicts the drug interaction is synergistic and a score >1 indicates an antagonistic drug interaction. We calculated MLSynergy scores for all two- and three-drug combinations of 8 frontline drugs (**File S3)**. We compared MLSynergy predictions of 26 two-drug and 40 three-drug combinations of the 8 frontline drugs to their DiaMOND assay-determined FIC_2_ and FIC_3_ scores, respectively (unpublished data from Bree Aldridge)^23,24^. This comparative analysis demonstrated that MLSynergy was >90% accurate in predicting synergistic and antagonistic effects of 2- and 3-drug combinations (**Figure 4D** and **4E**). Moreover, MLSynergy scores were highly correlated with the fractional inhibitory concentration values from DiaMOND (ρ=0.61, *p*-value < 0.001, **Figure S2**). Interestingly, three two-drug combinations (identified with red font in **Figure 4D**) were predicted as synergistic by MLSynergy, but were determined to be antagonistic by DiaMOND assay. Notably, these combinations were previously determined to be synergistic by other studies^23,25,26^, suggesting the effect of their drug interaction could be highly dependent on assay method and conditions. In contrast, MLSynergy is robust to the context in which a drug effect is measured since it was built on a SC-SVM classifier of Mtb viability from diverse growth conditions (**File S1**).

**Figure 4.**
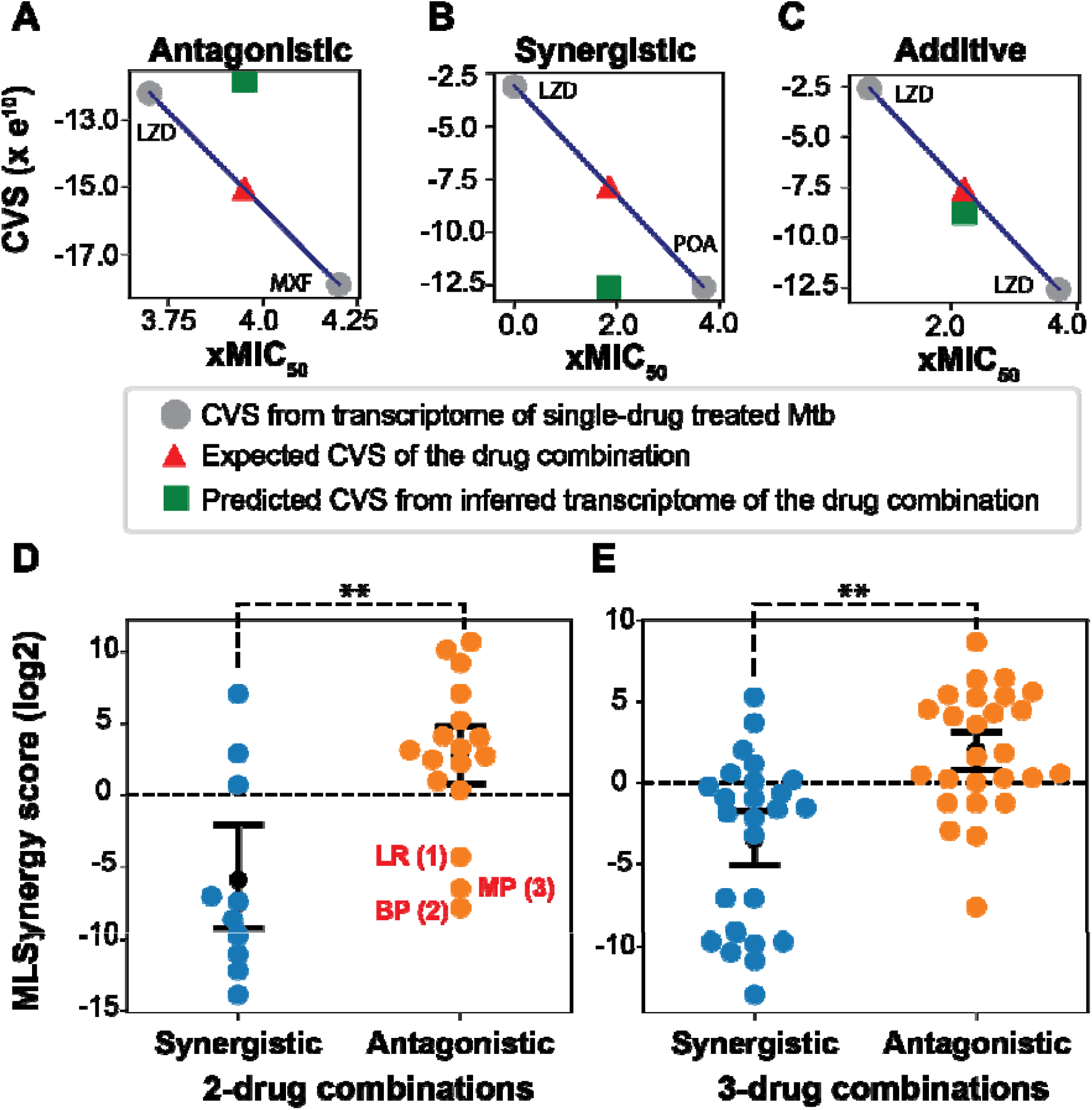
MLSynergy prediction of drug interaction. Examples of the relationship between expected CVS and predicted CVS for (**A**) antagonistic, (**B**) synergistic, and (**C**) additive drug combinations. The expected CVS (red triangle) was calculated as the average of DRonA-generated CVSs for experimentally measured transcriptomes from single-drug treated Mtb. LZD; linezolid, POA; pyrazinoic acid, MXF; moxifloxacin. (**D**) MLSynergy classification of experimentally validated synergistic and antagonistic 2-drug combinations. Drug combinations labeled in red: (1) LR; linezolid and rifampicin^25^, (2) BP; bedaquiline and pretomanid^23^, and (3) MP; moxifloxacin and pretomanid^38^ were classified as synergistic by MLSynergy and literature but antagonistic by DiaMOND assay. (**E**) MLSynergy classification of experimentally validated synergistic and antagonistic 3-drug combinations. Black dot and error bars indicate the mean and standard deviation away from the mean. Statistical significance (black dashed line) was calculated as *p*-value with Student’s T-test. **: *p*-value < 0.01.

## Discussion

Here, we report the first machine learning framework for drug response prediction in Mtb. DRonA enables efficient prediction of cell viability from transcriptomic signatures of perturbation, including drug treatment. Using DRonA estimates of cell viability from single-drug transcriptomic data, MLSynergy can then predict synergy and antagonism of antitubercular drug combinations. Our analysis using DRonA found a strong association between *in silico* estimates of cell viability following drug treatment and experimentally observed reduction in CFUs. Moreover, the loss of viability captured by DRonA from Mtb transcriptomes of patients undergoing HRZE treatment supports the clinical utility of our approach. Finally, we report validated and novel synergistic drug combinations, suggesting that the DRonA/MLSynergy framework is a promising tool for the prioritization of new multicomponent drug regimens. While we validated predictions of 2- and 3- drug interactions, our framework is generalizable for higher-order combinations.

The suitability of using the transcriptome as a reflection of Mtb viability was studied by treating Mtb with 7 frontline drugs and isolating RNA for transcriptome profiling while also evaluating cell viability by CFU. The DRonA predicted CVS of Mtb exposed to bactericidal (i.e., >MIC_50_) concentrations of drugs were below the cell viability threshold, proportional to relative CFU and significantly different to the CVS of untreated Mtb cultured for the same duration as drug treatment. Moreover, DRonA was able to perform effectively with other transcriptomic datasets of Mtb drug treatment, including during macrophage infection and from TB patients. The ability of DRonA to accurately predict the consequence of drug treatment in 7H9 medium, within macrophages, and from patient sputum, demonstrates that the definitions of viability in the DRonA model is inclusive of both actively dividing and slow replicating (physiologically adapted) phenotypes of Mtb. Moreover, the accuracy across datasets offers DRonA as a generalizable tool for use across drug response screens and in studies where gene expression was analyzed but Mtb viability was not measured.

Here we showed that DRonA complements bacteriological assays in evaluating treatment response. The decline in CVS corresponded to the decline in the proportions of surviving bacilli upon drug treatment, as measured by the relative CFU counts. Since no culturing is required, DRonA can estimate drug effects much faster than conventional bacteriological assays. Additionally, the ability to enrich and amplify RNA should allow DRonA to be used with samples where bacterial cell numbers are below the limit of detection of bacteriological assays. The high sensitivity and the autonomy from culturing, makes DRonA especially suitable to evaluate the efficacy of treatment regimens on drug tolerant Mtb subpopulations, such as persister cells ^27^.

Using DRonA-predicted viability scores, MLSynergy accurately predicted synergy and antagony for 2- and 3-drug combinations. This performance compares to INDIGO-MTB^28^, an existing strategy that quantifies synergistic and antagonistic drug regimens using transcriptomes of Mtb treated with individual drugs, but only with drugs with known drug-drug interactions. INDIGO-MTB requires known drug-drug interactions to learn patterns and identify combinations most likely to be synergistic. In contrast, the DRonA/MLSynergy platform is based on gene signatures of cell viability and does not require any input data related to drug combinations. Comparing the accuracy for drugs without prior drug interaction information, MLSynergy significantly outperforms INDIGO-MTB (*p*-value > 0.05, **Figure S3**). As such, our models can be more easily applied (i.e., no re-training required) to predict drug interaction for new drugs and conditions.

The DRonA/MLSynergy platform does have some potential limitations. First, our approach to predict cell viability based on gene expression signatures does not reveal information about drug mechanism of action. As cell death can lead to transcriptomic changes unrelated to drug treatment, these cell death signatures could be a confounding factor to make inferences about drug mechanism of action. Removing genes highly correlated to cell death could improve mechanism of action identification ^29^. Future work also aims to integrate DRonA with regulatory-metabolic networks ^30–33^ to reveal the underlying pathways that contribute to a drug’s bactericidal activity and interaction with other drugs.

A second limitation is that the DRonA/MLSynergy platform requires transcriptome profiling of Mtb drug treatment. It could be argued that testing viability is simpler and cheaper than transcriptomics. Yet, even for a small number of drugs, the resources needed to test the number of possible combinations quickly exceeds the cost of transcriptomics. Additionally, this is unlikely to be true when considering the facilities cost, maintenance and training required for bacteriological assays with Mtb. The cost of transcriptomes will be less of a concern in the future as transcriptomes are increasingly becoming faster and less costly to generate. New instruments and methods are also enabling transcriptomes directly from patient samples ^8^, which could permit DRonA as a rapid point-of-care evaluation of the effectiveness of drug treatment in TB patients.

Drug response prediction with machine learning models is an important area of current research, particularly for a slow-growing pathogen, and our results highlight the practicality of using transcriptome signatures to address major bottlenecks in the drug discovery process. The ability to detect changes in cell viability and predict drug interaction using just transcriptome profiles could substantially accelerate TB drug discovery efforts. Finally, the DronA/MLSynergy framework can be easily extended to predict other phenotypes of Mtb associated with gain of drug resistance (e.g., metabolic states, and cell wall composition), which could further improve treatment response prediction and clinical outcomes. To facilitate the widespread usage of these resources, the compendia of Mtb transcriptomes collated from GEO, and generated in this study have been deposited in GEO, and are publically available together with DRonA and MLSynergy algorithms and models in GitHub (https://github.com/baliga-lab/DRonA_MLSynergy), and in the Mtb Network portal (http://networks.systemsbiology.net/mtb/).

## Acknowledgements

We thank Amardeep Kaur for her technical expertise; Mario L. Arrieta-Ortiz and Adrian Garcia Lopez de Lomana for help during the development of DRonA and MLSynergy; and other members of the Baliga lab for critical discussions. Funding was provided by the Bill and Melinda Gates Foundation (INV-009322) and the National Institute of Allergy and Infectious Diseases (R01AI128215, R01AI141953 and U19AI135976).

## Author contributions

V.S. designed research, wrote GEOparser, DRonA and MLSynergy algorithms, performed drug assays, analyzed data and wrote the paper. R.A.R. performed drug assays and collected samples for transcriptomics. M.P. prepared the RNA seq libraries. S.R.C.I aided in algorithm development. E.J.R.P and N.S.B. designed research, analyzed data, and wrote the paper.

## Competing interests

The authors declare no competing financial interests

## Material and methods

### Bacterial strains and growth conditions

*Mycobacterium tuberculosis* strains used in this study was H37Rv. Mtb cells were culture in standard 7H9-rich media consisting of 7H9 broth with 0.05% Tween-80, 0.2% glycerol, and 10% Middlebrook ADC. Frozen 1 mL stocks of Mtb cells were added to 7H9 medium and grown with mild agitation in a 37° C incubator until the culture reached an OD_600_ of ∼0.4-0.8. The cells were then diluted to OD_600_ of 0.05 and added to 7H9-rich medium containing drugs at the predetermined amounts.

### Minimum Inhibitory Concentration 50 (MIC_50_) determination

10 mM working concentrations of drugs considered in the study were made with a suitable vehicle depending on drug solubility (*i*.*e*., water, DMSO, or methanol). The 10 mM working concentrations of drugs were diluted in a twofold dilution series for 11 concentrations in 96-well plates. Each treatment series contained an untreated well as a control. Mtb H37Rv cultures were added to the wells and the plates were incubated at 37° C. Growth in cultures were measured as OD_600_ at 0 and 72 hours of incubation. All MIC_50_ determinations were performed in biological triplicate. Growth inhibition was determined by subtracting the initial reads from the final reads and then normalizing the data to no drug controls. Growth inhibition was fit to a sigmoidal curve and MIC_50_ was calculated for each drug (**Table S1**).

### Time-kill assays

Using growth conditions described above, cells were diluted into 7H9-rich media containing drugs at predetermined amounts, along with vehicle controls (**Table S2**). Samples were taken after 0, 24 and 72 h, serially diluted and plated on 7H10 agar plates. All time-kill assays were performed in biological triplicate. Relative CFUs were calculated as log_10_ ratio of CFUs/ml of culture observed at start of treatment (T_0_) and after drug treatment.

### Collection, RNA extraction, and analysis of single-drug transcriptomes

Using growth conditions described above, cells were diluted into 7H9-rich media containing drugs at predetermined amounts, along with vehicle controls (**Table S1** and **File S2**). Samples, in biological triplicates, were collected after 24 and 72 h. Samples were centrifuged at high speed for 5 min, supernatant was discarded and cell pellet was immediately flash frozen in liquid nitrogen. Cell pellets were stored at -80° C until bead beating in a FastPrep 120 homogenizer and RNA extraction was performed as previously described ^34^. Total RNA was depleted of ribosomal RNA using the Ribo-Zero Bacteria rRNA Removal Kit (Illumina, San Diego, CA). Quality and purity of the mRNA was determined with 2100 Bioanalyzer (Agilent, Santa Clara, CA). Sequencing libraries were prepared with TrueSeq Stranded mRNA HT library preparation kit (Illumina, San Diego, CA). All samples were sequenced on the NextSeq sequencing instrument in a high output 150 v2 flow cell. Paired-end 75 bp reads were checked for technical artifacts using Illumina default quality filtering steps. Raw FASTQ read data were processed using the R package DuffyNGS as previously described^35^. Read counts were further analyzed with Kallisto^36^ and RPKM values were calculated.

### Collection and curation of Mtb-GEO dataset for training of DronA

GEOParser was developed to download transcript profiles and metadata of drug-treated and untreated samples of Mtb-H37Rv from Gene Expression Omnibus (GEO). GEOparser collected median spot intensity from microarray samples and Reads Per Kilobase of transcript, per Million mapped reads (RPKM) from RNA-seq samples. The compendium dataset was curated by removing samples with low coverage (i.e., samples with <70% of annotated Mtb genes). The curated dataset was normalized by rank normalization with the “minimum” method with SciPy package^37^. Using the metadata collected by GEOParser, manual classification determined the transcriptomes belonging to the “viable” class used to seed DRonA and the “drug treatment” class for testing (**File S1**).

### Training and running DronA

The initial training set of manually classified “viable” transcriptomes (**File S1**) was used to start the iterative training of DRonA. Each iteration consisted of the following steps: (1) a single class support vector machine (SC-SVM) was trained on the training set; (2) the accuracy of the trained SC-SVM was calculated with the test set; (3) assessment of the accuracy, and (4) transcriptomes from the non-classified set that classified as viable were identified and added to the training set. The iterative process was stopped when the accuracy of the classifier dropped below a false positive rate threshold (80%) or when no new transcriptomes from non-classified set were added to the training set. The cell viability scores (CVS) were calculated for samples as the weighted sum of gene expression ranks using the trained SC-SVM. CVSs were normalized by subtracting the score of a sample with maximum score observed in that experiment.

### Inference of multi-drug transcriptomes

Transcriptomes of the Mtb cultures treated with multi-drug combinations at effective doses were predicted by triangulation with the single-drug treated transcriptomes and untreated control. Triangulation was called through ‘triangulate’ function in the MLSynergy algorithm, which collects transcriptomes of the drugs in combination (each profiled as single-drug) and untreated control and averages them with geometric mean. The inferred multi-drug transcriptomes were then returned to DRonA for CVS determination.

### Calculation of MLSynergy scores for drug combinations

CVSs were obtained from DRonA with the transcriptomes of the single-drug treatments that make up the drug combination and expected CVS was calculated by averaging the CVSs of single-drug treatments. The predicted CVS was obtained from DRonA with the inferred transcriptome of the drug combination. MLSynergy score were calculated as the ratio of expected CVS and predicted CVS. Further, MLSynergy scores were log normalized (base 2) in reference to the average of MLSynergy scores of same drug combinations that are considered to be additive in nature.

### Comparison of INDIGO-MTB and MLSynergy predictions

Two INDIGO^28^ models were retrained with default parameters. Model-1 was trained with the complete dataset (202 combinations and 46 drugs) and Model-2 was trained with partial dataset (98 combinations and 40 drugs) which was obtained after excluding combinations with bedaquiline, clofazimine, linezolid, moxifloxacin, pretomanid and pyrazinamide. Both models were tested on the combinations given in **File S4**. Transcriptomes provided in Ma *et al*. were used as input for the INDIGO models. Transcriptomes generated in this study (summarized in **File S3**) were used as input for the MLSynergy.

## Code and data availability

Codes for GEOparser, DRonA and MLSynergy and the training data used to train DRonA is available in the GitHub (https://github.com/baliga-lab/DRonA_MLSynergy). Transcriptomes generated as part of this study are available in the gene expression omnibus database (GEO accession number: GSE16567).

## Abbreviations

Mtb: *Mycobacterium tuberculosis*
TB: Tuberculosis
MIC: Minimum inhibitory concentration
GEO: Gene expression omnibus
SC-SVM: Single class support vector machine
CVS: Cell viability score
CFU: Colony forming unit
DRonA: Drug Response Assayer

## Supplementary materials

**Table S1.**
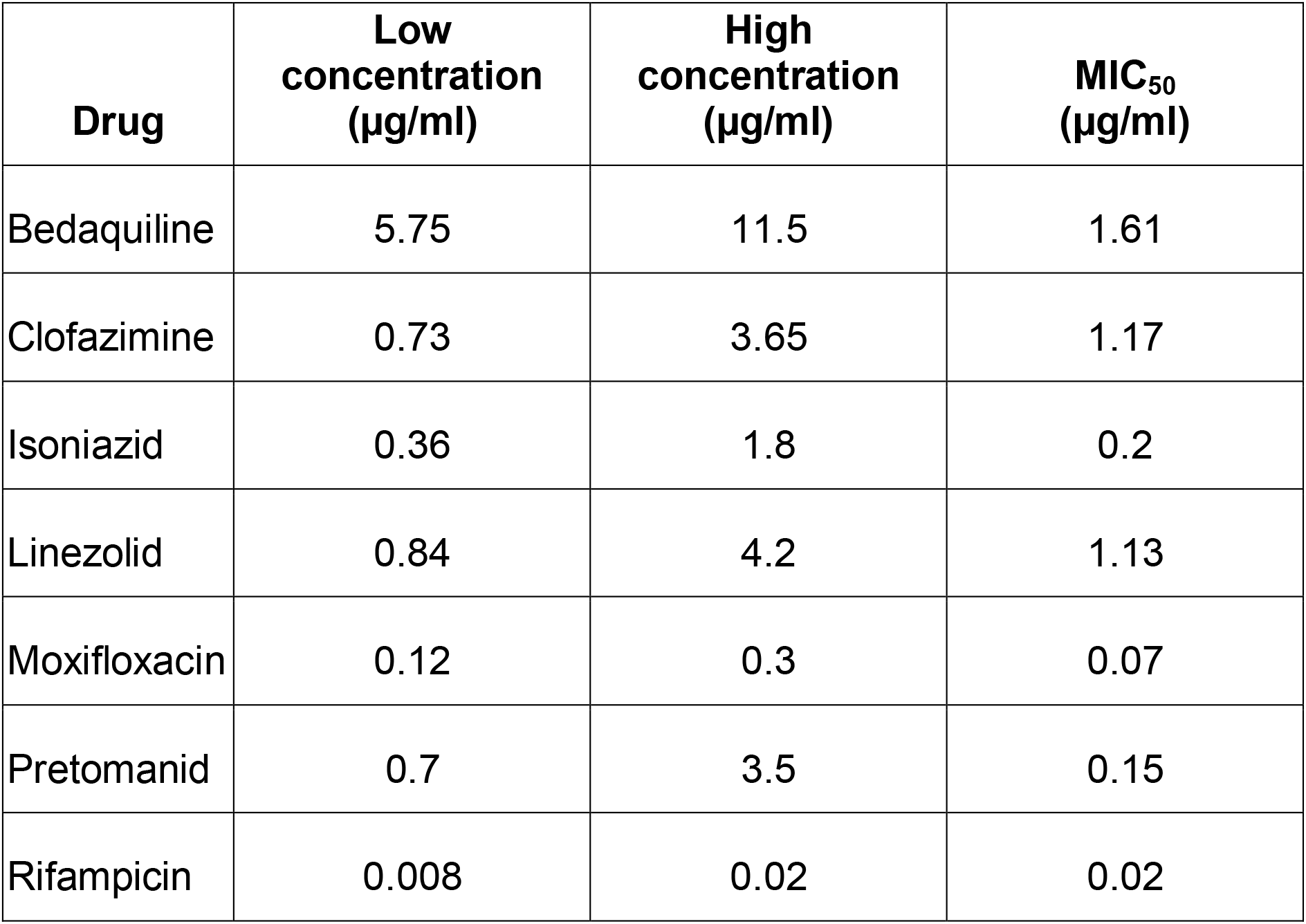
MIC_50_ determination of the drugs used in the study and drug concentrations used for transcriptome profiles generated in this study.

**Table S2.** Drug treatment concentrations and time used to compare the DRonA generated CVS and relative CFUs (related to **Figure 3**).

| Legend | Drug, Concentration (µg/ml), Treatment time in hours |
| --- | --- |
| 1 | BDQ, 5.75, 24 |
| 2 | BDQ, 5.75, 72 |
| 3 | CFZ, 0.73, 24 |
| 4 | CFZ, 0.73, 72 |
| 5 | INH, 0.36, 24 |
| 6 | INH, 0.36, 72 |
| 7 | INH, 1.80, 24 |
| 8 | INH, 1.80, 72 |
| 9 | LZD, 0.84, 24 |
| 10 | LZD, 0.84, 72 |
| 11 | MXF, 0.12, 24 |
| 12 | MXF, 0.12, 72 |
| 13 | MXF, 0.30, 24 |
| 14 | MXF, 0.30, 72 |
| 15 | No drug, 0.00, 24 |
| 16 | No drug, 0.00, 72 |
| 17 | PA824, 0.70, 24 |
| 18 | PA824, 0.70, 72 |
| 19 | RIF, 0.01, 24 |
| 20 | RIF, 0.01, 72 |
| 21 | RIF, 0.02, 24 |
| 22 | RIF, 0.02, 72 |

**Figure S1.**
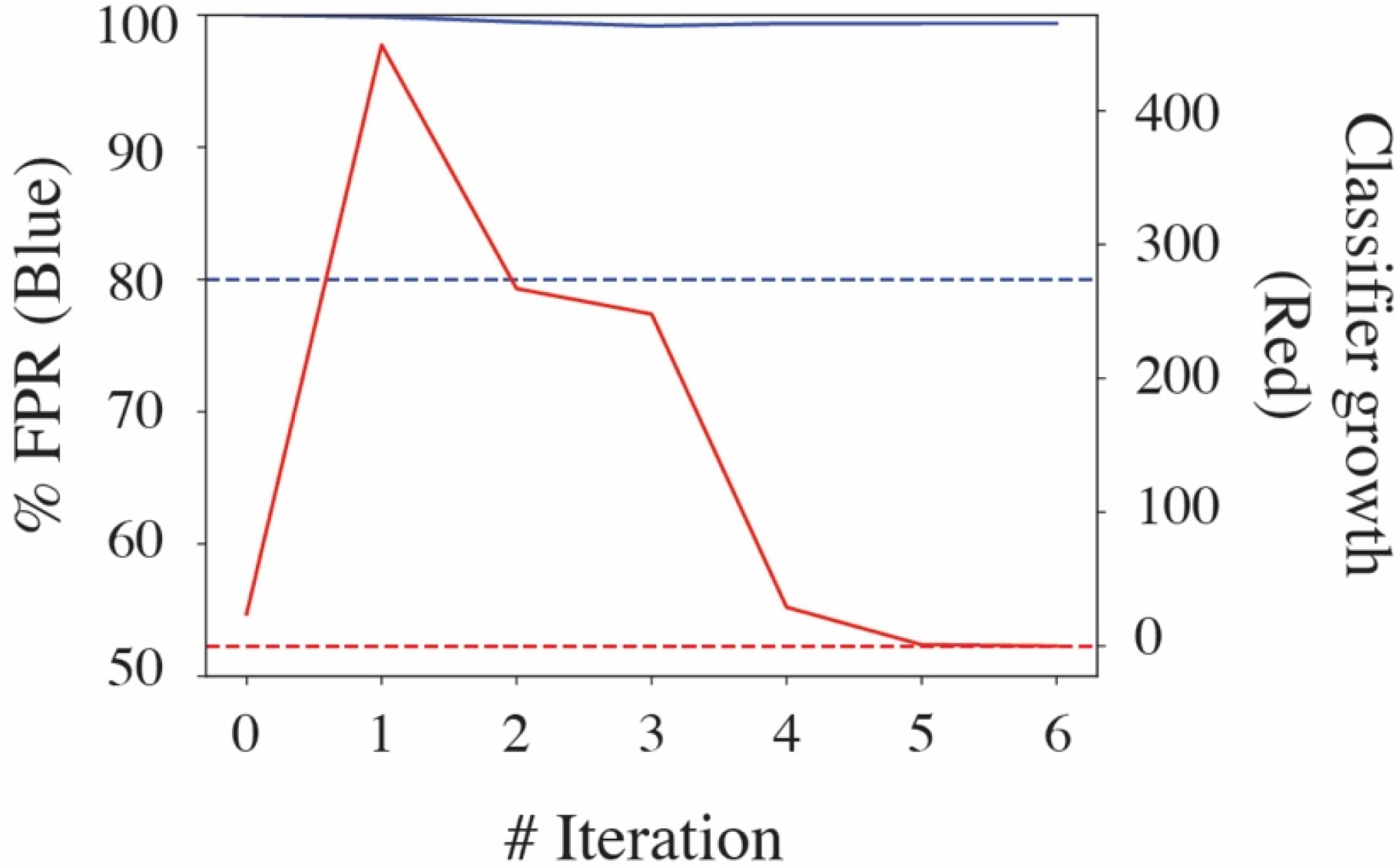
Iterative training of DRonA. The graph shows changes in accuracy, measured as percentage false positive rate (% FPR in Blue) and number of non-classified transcriptomes added to the viable training (Classifier growth in Red) at each iteration. The % FPR was calculated as FP/N, where FP is the number of drug treated non-viable transcriptomes (from test set) that were classified as viable and N is the total number of non-viable transcriptomes (from test set). Blue and red dashed line show the threshold for % FPR and classifier growth below which the iterative training was programmed to stop. The classifier growth dropped below threshold at iteration #5 and was halted.

**Figure S2.**
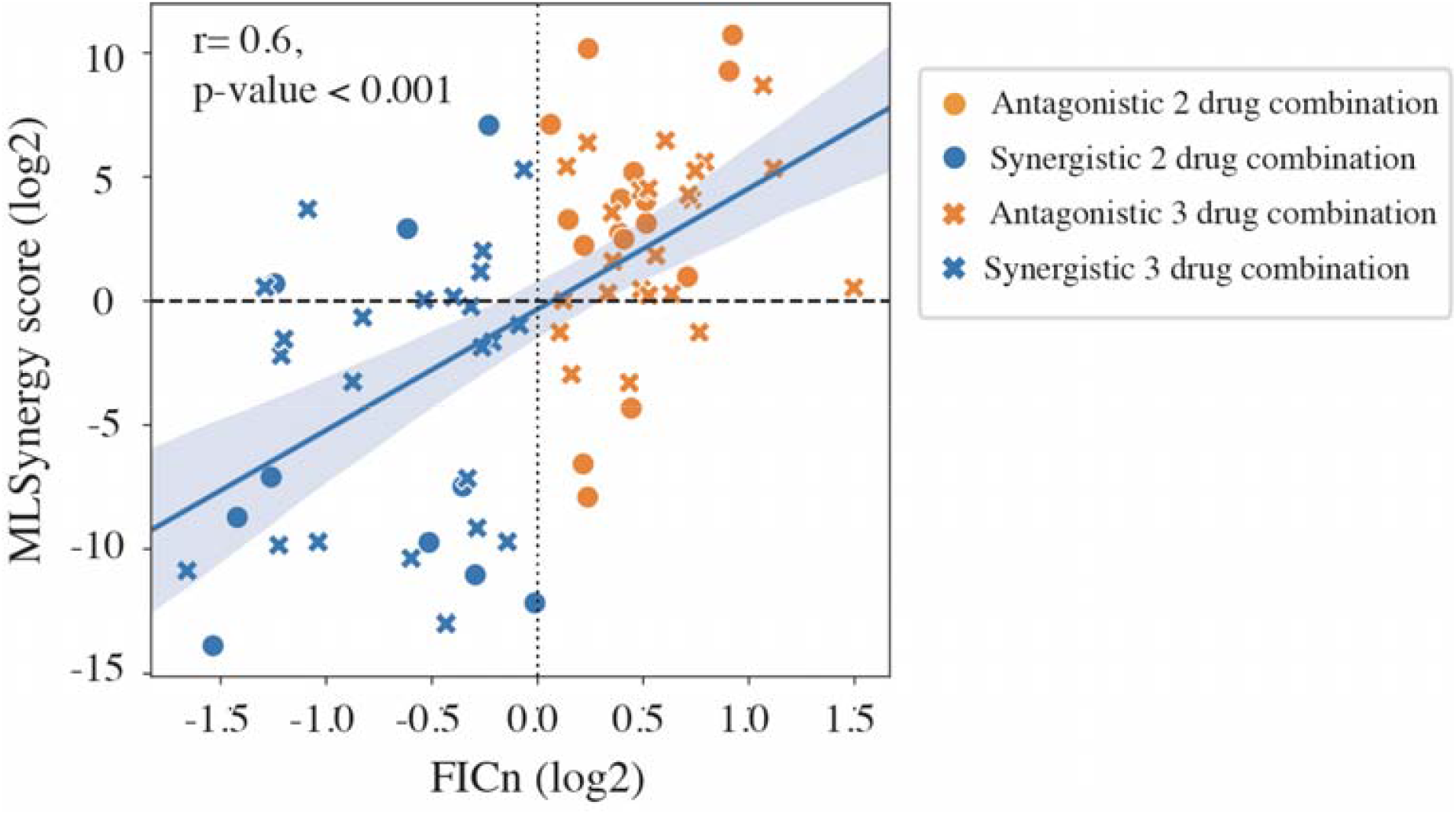
Correlation between MLSynergy score and FIC_n_ ^23,24^ for 2- and 3-drug combinations.. Solid blue line denotes the Pearson’s correlation between CVS and relative CFU. Significance was calculated as the average correlation coefficient, *r*, from 100 iterations performed with 70% randomly selected data. Black dotted line and black dashed line are the FIC_n_ score and MLSynergy scores, respectively, that separate synergistic combinations from the antagonistic combinations.

**Figure S3.**
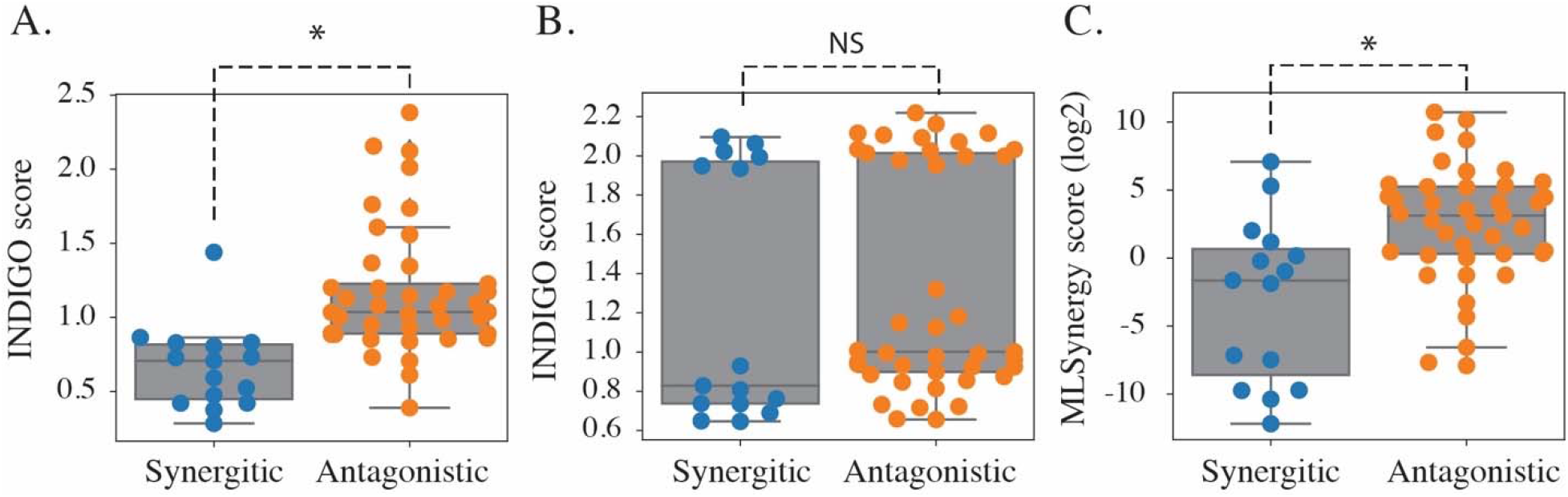
Comparison of INDIGO^28^ models with MLSynergy in predicting interaction of 2- and 3-drug combinations. (**A**) Predictions from INDIGO model that was trained on 202 drug combinations with 46 drugs (Model-1) (**B**) Predictions from INDIGO model that was trained on 98 drug combination with 40 drugs (Model-2); specifically combinations with bedaquiline, clofazimine, linezolid, moxifloxacin, pretomanid and pyrazinamide were excluded from training, and (**C**) Predictions from MLSynergy model with same drug combinations as (**B**). Drugs were validated as synergistic and antagonistic from DiaMOND assay. Data plotted is tabulated in **File S4**.

---

**File S1**. Metadata of the GEO compendium used for training DRonA.

**File S2**. (A) Transcriptome profiles generated in this study to test DRonA, related to Figure 2A. (B) Transcriptomes from Liu *et al*. used to test DRonA, related to Figure 2B.

**File S3**. MLSynergy interaction prediction for 2- and 3-drug combinations. Drug concentrations given as the factor of MIC_50_s from Table S1. BDQ: bedaquiline, CFZ: clofazimine, INH: isoniazid, LZD: linezolid, MXF: moxifloxacin, PA824: pretomanid, RIF: rifampicin, POA: pyrazinoic aicd.

**File S4**. INDIGO (both models) and MLSynergy scores for 2- and 3-drug combinations plotted in Figure S4.

## Notes

### Competing Interest Statement

The authors have declared no competing interest.

### Summary of Updates

Authors were linked with their ORCIDs.

## Reference

1. Ji, B., Levy, L. & Grosset, J. H. Chemotherapy of leprosy: progress since the Orlando Congress, and prospects for the future. Int. J. Lepr. Mycobact. Dis. Off. Organ Int. Lepr. Assoc. 64, S80-88; discussion S88-90 (1996).

2. Sarathy, J. et al. Fluoroquinolone Efficacy against Tuberculosis Is Driven by Penetration into Lesions and Activity against Resident Bacterial Populations. Antimicrob. Agents Chemother. 63, (2019).

3. Sarathy, J. P. et al. Extreme Drug Tolerance of Mycobacterium tuberculosis in Caseum. Antimicrob. Agents Chemother. 62, (2018).

4. Diacon, A. H. et al. Time to positivity in liquid culture predicts colony forming unit counts of Mycobacterium tuberculosis in sputum specimens. Tuberc. Edinb. Scotl. 94, 148–151 (2014).

5. Honeyborne, I. et al. Molecular bacterial load assay, a culture-free biomarker for rapid and accurate quantification of sputum Mycobacterium tuberculosis bacillary load during treatment. J. Clin. Microbiol. 49, 3905–3911 (2011).

6. Honeyborne, I. et al. The molecular bacterial load assay replaces solid culture for measuring early bactericidal response to antituberculosis treatment. J. Clin. Microbiol. 52, 3064–3067 (2014).

7. Honeyborne, I. et al. Profiling persistent tubercule bacilli from patient sputa during therapy predicts early drug efficacy. BMC Med. 14, 68 (2016).

8. Peterson, E. J. et al. Path-seq identifies an essential mycolate remodeling program for mycobacterial host adaptation. Mol. Syst. Biol. 15, e8584 (2019).

9. Walter, N. D. et al. Transcriptional Adaptation of Drug-tolerant Mycobacterium tuberculosis During Treatment of Human Tuberculosis. J. Infect. Dis. 212, 990–998 (2015).

10. Peterson, N. D., Rosen, B. C., Dillon, N. A. & Baughn, A. D. Uncoupling Environmental pH and Intrabacterial Acidification from Pyrazinamide Susceptibility in Mycobacterium tuberculosis. Antimicrob. Agents Chemother. 59, 7320–7326 (2015).

11. Liu, Y. et al. Immune activation of the host cell induces drug tolerance in Mycobacterium tuberculosis both in vitro and in vivo. J. Exp. Med. 213, 809–825 (2016).

12. Andreu, N., Fletcher, T., Krishnan, N., Wiles, S. & Robertson, B. D. Rapid measurement of antituberculosis drug activity in vitro and in macrophages using bioluminescence. J. Antimicrob. Chemother. 67, 404–414 (2012).

13. Salfinger, M., Crowle, A. J. & Reller, L. B. Pyrazinamide and pyrazinoic acid activity against tubercle bacilli in cultured human macrophages and in the BACTEC system. J. Infect. Dis. 162, 201–207 (1990).

14. Adams, K. N. et al. Drug Tolerance in Replicating Mycobacteria Mediated by a Macrophage-Induced Efflux Mechanism. Cell 145, 39–53 (2011).

15. Fang, F. et al. LPS restores protective immunity in macrophages against Mycobacterium tuberculosis via autophagy. Mol. Immunol. 124, 18–24 (2020).

16. Koul, A. et al. Delayed bactericidal response of Mycobacterium tuberculosis to bedaquiline involves remodelling of bacterial metabolism. Nat. Commun. 5, (2014).

17. Koul, A. et al. Diarylquinolines are bactericidal for dormant mycobacteria as a result of disturbed ATP homeostasis. J. Biol. Chem. 283, 25273–25280 (2008).

18. Koul, A. et al. Diarylquinolines target subunit c of mycobacterial ATP synthase. Nat. Chem. Biol. 3, 323–324 (2007).

19. Andries, K. et al. A diarylquinoline drug active on the ATP synthase of Mycobacterium tuberculosis. Science 307, 223–227 (2005).

20. Peterson, E. J. R., Ma, S., Sherman, D. R. & Baliga, N. S. Network analysis identifies Rv0324 and Rv0880 as regulators of bedaquiline tolerance in Mycobacterium tuberculosis. Nat. Microbiol. 1, microbiol201678 (2016).

21. Loewe, S. Die quantitativen Probleme der Pharmakologie. Ergeb. Physiol. 27, 47–187 (1928).

22. Deshpande, D. et al. Concentration-Dependent Synergy and Antagonism of Linezolid and Moxifloxacin in the Treatment of Childhood Tuberculosis: The Dynamic Duo. Clin. Infect. Dis. Off. Publ. Infect. Dis. Soc. Am. 63, S88–S94 (2016).

23. Cokol, M., Kuru, N., Bicak, E., Larkins-Ford, J. & Aldridge, B. B. Efficient measurement and factorization of high-order drug interactions in Mycobacterium tuberculosis. Sci. Adv. 3, (2017).

24. Larkins-Ford, J. et al. Systematic measurement of combination drug landscapes to predict in vivo treatment outcomes for tuberculosis. bioRxiv 2021.02.03.429579 (2021) doi:10.1101/2021.02.03.429579.

25. Diacon, A. H. et al. 14-day bactericidal activity of PA-824, bedaquiline, pyrazinamide, and moxifloxacin combinations: a randomised trial. Lancet Lond. Engl. 380, 986–993 (2012).

26. Maltempe, F. G. et al. Activity of rifampicin and linezolid combination in Mycobacterium tuberculosis. Tuberculosis 104, 24–29 (2017).

27. Srinivas, V., Arrieta-Ortiz, M. L., Kaur, A., Peterson, E. J. R. & Baliga, N. S. PerSort Facilitates Characterization and Elimination of Persister Subpopulation in Mycobacteria. mSystems 5, (2020).

28. Ma, S. et al. Transcriptomic Signatures Predict Regulators of Drug Synergy and Clinical Regimen Efficacy against Tuberculosis. mBio 10, (2019).

29. Szalai, B. et al. Signatures of cell death and proliferation in perturbation transcriptomics data—from confounding factor to effective prediction. Nucleic Acids Res. 47, 10010–10026 (2019).

30. Turkarslan, S. et al. A comprehensive map of genome-wide gene regulation in Mycobacterium tuberculosis. Sci. Data 2, 150010 (2015).

31. Peterson, E. J. R. et al. A high-resolution network model for global gene regulation in Mycobacterium tuberculosis. Nucleic Acids Res. 42, 11291–11303 (2014).

32. Chandrasekaran, S. & Price, N. D. Probabilistic integrative modeling of genome-scale metabolic and regulatory networks in Escherichia coli and Mycobacterium tuberculosis. Proc. Natl. Acad. Sci. 107, 17845–17850 (2010).

33. Immanuel, S. R. C. et al. Quantitative prediction of conditional vulnerabilities in regulatory and metabolic networks of Mycobacterium tuberculosis. bioRxiv 2021.01.29.428876 (2021) doi:10.1101/2021.01.29.428876.

34. Peterson, E. J. R. et al. Intricate Genetic Programs Controlling Dormancy in Mycobacterium tuberculosis. Cell Rep. 31, 107577 (2020).

35. Vignali, M. et al. NSR-seq transcriptional profiling enables identification of a gene signature of Plasmodium falciparum parasites infecting children. J. Clin. Invest. 121, 1119– 1129 (2011).

36. Bray, N. L., Pimentel, H., Melsted, P. & Pachter, L. Near-optimal probabilistic RNA-seq quantification. Nat. Biotechnol. 34, 525–527 (2016).

37. Virtanen, P. et al. SciPy 1.0: fundamental algorithms for scientific computing in Python. Nat. Methods 17, 261–272 (2020).

38. de Miranda Silva, C. et al. Effect of Moxifloxacin plus Pretomanid against Mycobacterium tuberculosis in Log Phase, Acid Phase, and Nonreplicating-Persister Phase in an In Vitro Assay. Antimicrob. Agents Chemother. 63, (2019).

